# Prospective Study of Polygenic Risk, Protective Factors, and Incident Depression Following Combat Deployment in US Army Soldiers

**DOI:** 10.1101/361725

**Authors:** Karmel W. Choi, Chia-Yen Chen, Robert J. Ursano, Xiaoying Sun, Sonia Jain, Ronald C. Kessler, Karestan C. Koenen, Min-Jung Wang, Gary H. Wynn, Major Depressive Disorder Working Group of the Psychiatric Genomics Consortium, Laura Campbell-Sills, Murray B. Stein, Jordan W. Smoller

## Abstract

**Background:** Whereas genetic susceptibility increases risk for major depressive disorder (MDD), non-genetic protective factors may mitigate this risk. In a large-scale prospective study of US Army soldiers, we examined whether trait resilience and/or unit cohesion could buffer against the onset of MDD following combat deployment, even in soldiers at high polygenic risk.

**Methods:** Data were analyzed from 4,182 soldiers of European ancestry assessed before and after their deployment to Afghanistan. Incident MDD was defined as no MDD episode at predeployment, followed by a MDD episode following deployment. Polygenic risk scores were constructed from the largest available MDD genome-wide association study. We first examined main effects of the MDD PRS and each protective factor on incident MDD. We then tested effects of each protective factor on incident MDD across strata of polygenic risk.

**Results:** Polygenic risk showed a dose-response relationship to depression, such that soldiers at high polygenic risk had greatest odds for incident MDD. Both unit cohesion and trait resilience were prospectively associated with reduced risk for incident MDD. Notably, the protective effect of unit cohesion persisted even in soldiers at highest polygenic risk.

**Conclusions:** Polygenic risk was associated with new-onset MDD in deployed soldiers. However, unit cohesion—an index of perceived support and morale—was protective against incident MDD even among those at highest genetic risk, and may represent a potent target for promoting resilience in vulnerable soldiers. Findings illustrate the value of combining genomic and environmental data in a prospective design to identify robust protective factors for mental health.

## Introduction

Exposure to stressful experiences is an important risk factor for major depressive disorder (MDD) (1)—however, not all individuals exposed to stressful experiences develop MDD. This observation is of high relevance for the US Army, whose soldiers routinely encounter stressful events over the course of combat deployment (2) and show a correspondingly high burden of MDD following deployment (3–5). Preventing MDD and its associated disability and comorbidities can improve individual/family wellbeing and troop readiness (6), and requires attention to risk and protective factors that influence depression.

The diathesis-stress model of depression (7) posits that some individuals have latent or pre-existing vulnerabilities, or *diatheses*, that are activated in the presence of stress to produce MDD. One such diathesis is genetic susceptibility, which has been found to substantially increase risk for MDD episodes in the presence of stressful life events (8). Genetic susceptibility for a complex trait like MDD is thought to be polygenic—influenced by many common variants across the genome, each with relatively small effect sizes (9). This influence can be indexed by polygenic risk scores (PRS) that combine effects across common variants using results from a discovery genome-wide association study (GWAS). For MDD, a well-powered GWAS with 461,134 individuals (10) has become available, and its derived PRS was recently validated in a diathesis-stress model for MDD in the context of life stressors (11). However, to date, this PRS has not been prospectively validated in terms of new MDD onset following stress exposure.

While genetic susceptibility is a risk factor for depression, non-genetic protective factors may buffer this risk, illuminating opportunities for prevention. Protective factors can be specific to the individual (intrinsic) or related to the individual’s environment (extrinsic) (12). One intrinsic factor that has been studied in Army populations is trait resilience, defined as perceived hardiness to stress and ability to cope adaptively with stressors (13). Unit cohesion—which includes emotional safety, bonding, and support between soldiers and with unit leaders—is an extrinsic factor that has also received substantial attention. Although these factors are well characterized for their protective effects on post-deployment mental health (14–17), the extent to which they attenuate risk for MDD in the presence of genetic susceptibility has not been examined.

The Army Study of Risk and Resilience in Servicemembers (Army STARRS) has followed a large prospective sample of active duty soldiers with genomic data across one combat deployment cycle (18). This provides a unique opportunity to test the effects of genetic susceptibility and candidate protective factors assessed shortly before deployment, in relation to development of MDD following deployment. Specifically, we examine whether two putative protective factors—trait resilience (intrinsic) and unit cohesion (extrinsic)—can reduce risk for incident post-deployment MDD even among soldiers at high polygenic risk for MDD.

## Methods

### Participants and procedures

The Pre/Post Deployment Study (PPDS) in Army STARRS is a multi-wave panel survey of US Army soldiers from three brigade combat teams that were deployed to Afghanistan in 2012. Soldiers completed baseline assessments within approximately six weeks before deployment, and follow-up assessments at three and nine months post-deployment. For this analysis, the sample was restricted to those with eligible survey responses and samples for genotyping (N=4,900). Procedures for Army STARRS and PPDS have been reported in detail elsewhere (18,19). All participating soldiers provided written informed consent, and study procedures were approved by the institutional review boards at the Uniformed Services University of the Health Sciences, Harvard University, University of Michigan, and University of California, San Diego.

### Measures

#### Major depressive disorder (MDD)

MDD was ascertained at each assessment using items from the major depressive episode (MDE) scale of the WHO Composite International Diagnostic Interview-Screening Scales (CIDI-SC) (20). Scale items assessed frequency of MDD symptoms (e.g., depressed mood, loss of interest) over the past 30 days, and were summed to yield overall symptom scores. Symptom scores were then dichotomized using receiver operating characteristic (ROC) curve analysis to determine clinical thresholds for past 30-day MDEs, as validated elsewhere (20). Incident MDD was defined as no MDE at baseline, followed by a MDE at any point through nine months (0=no incident depression, 1=incident depression). Soldiers who met criteria for an existing MDE at pre-deployment (N=310) or had no follow-up MDE data (N=408) were excluded since MDD incidence could not be established.

#### Trait resilience

Trait resilience was self-reported by soldiers at baseline using a five-item scale derived from a larger pool of 17 items that were pilot-tested in earlier Army STARRS surveys and culled using exploratory factor analysis and item response theory analysis for administration in the PPDS. Information on the development and validation of this scale has been published elsewhere (21). Participants reported on their abilities to “keep calm and think of the right thing to do in a crisis,” “manage stress,” or to “try new approaches if old ones don’t work” (all items described in *Supplementary Materials S1A*). Items were rated on a five-point Likert scale ranging from “poor” to “excellent,” and summed to yield continuous scores ranging from 0 and 20. Internal consistency was good (α=0.89). Scores were standardized to a mean of 0 and variance of 1 for analysis.

#### Unit cohesion

Unit cohesion was assessed at baseline using a seven-item scale developed for this study and adapted from the Walter Reed Army Institute of Research (WRAIR) Military Cohesion Scales (22). Soldiers reported on perceived support and cohesion within their unit, including items such as “I can rely on members of my unit for help if I need it,” “I can open up and talk to my first line leaders if I need help,” and “My leaders take a personal interest in the well-being of all soldiers in my unit” (all items described in *Supplementary Materials S1B*).

Items were rated on five-point Likert scale ranging from “strongly disagree” to “strongly agree,” and were summed to yield continuous scores ranging between 0 and 27. Internal consistency was high (α=0.89); a factor analysis confirmed that one single factor was sufficient to represent shared variability among these seven scale items. Scores were standardized to a mean of 0 and variance of 1 for analysis.

#### Combat stress exposure

Within one month of return from deployment, soldiers also completed a 15-item measure of combat stress exposure—including engaging in combat patrol or other dangerous duties, and firing at and/or receiving enemy fire. These items were summed to reflect overall burden of combat stress exposure, as in previous research (19).

### DNA processing

Detailed information about genotyping, imputation, quality control (QC), and population assignment in Army STARRs is available elsewhere (23). Briefly, DNA samples for each participant were genotyped using Illumina OmniExpress and Exome array with additional custom content. Initial quality control (QC) procedures were conducted to retain only (1) samples with genotype missingness < 0.02, no extreme autosomal heterozygosity, and no relatedness (if related pairs of individuals were identified, only one was kept); and (2) single-nucleotide polymorphisms (SNPs) with genotype missingness < 0.05 (before sample QC) and < 0.02 (after sample QC), minor allele frequency (MAF) > 0.05, and no violation of the Hardy-Weinberg equilibrium (p > 1×10^−6^).

Prior to imputation, SNPs were also removed if they were not present or had non-matching alleles in the 1000 Genomes Project reference panel (24) or had ambiguous alleles with MAF > 0.10. Following a two-step pre-phasing/imputation process (25), imputed SNPs were converted to “best guess” genotyped SNPs based on their imputation probability. Where no possible genotype met the threshold of 80% probability, information for that SNP was set as missing. SNPs were filtered again to retain missingness < 0.02 and imputation quality (INFO) score > 0.80, and duplicate SNPs were identified for exclusion in subsequent analyses.

Ancestry was inferred through principal component (PC) analyses as reported previously (23). Given that polygenic risk scores would be constructed using effect sizes obtained in samples of European ancestry (10), only PPDS participants assigned to the European ancestry (EA) group were retained for this study. By inspecting successive PC plots within the EA group for evidence of population structure, we determined only the first three principal components were likely relevant for inclusion as covariates in subsequent analyses.

### Polygenic risk scoring

To construct the polygenic risk scores (PRS), we obtained summary statistics from the latest GWAS of MDD in 461,134 individuals (10). For main analyses, we used the set of summary statistics without 23andMe data (N=173,005) that is now publicly available. After removal of ambiguous SNPs, we clumped the GWAS summary statistics using our EA genomic data to limit inclusion of highly correlated SNPs, using a r^2^ threshold of 0.25 and a 250kb window. These clumped summary statistics were used to compute PRS from our EA genomic data that included SNPs whose effects met the following p-value thresholds (pT) in decreasing order of stringency: 5×10^−8^, <0.0001, <0.001, 0.01, 0.05, 0.10, 0.50, 1.0. PRS were calculated as the total sum of risk alleles at each eligible SNP weighted by their estimated effect size (log odds ratio), divided by total number of SNPs included for scoring (*Supplementary Table S2A*).

### Statistical analyses

First, we examined the MDD PRS at varying p-value thresholds in relation to incident MDD (*Supplementary Materials Figure S2A*). The PRS at the p-value threshold with largest Nagelkerke’s pseudo-R^2^ (pT=0.01) was selected for subsequent analyses (26). This PRS was distributed across individuals and divided into three groups of polygenic risk (*Supplementary Table S2B*): low (quintile 1), intermediate (quintiles 2-4), and high (quintile 5) (27). Second, we used logistic regressions to examine the main effects of polygenic risk on incident MDD, using the low risk group as the reference group. Third, we used logistic regressions to examine the main effects of each protective factor on incident MDD. Fourth, we tested the effects of each protective factor (per standardized unit score) on incident MDD across polygenic risk groups. At each step, we adjusted for sex, age, and principal components to account for population stratification when polygenic scores were included. All analyses were conducted in R.

## Results

### Sample characteristics

Our sample included PPDS participants of European ancestry who provided genome-wide data, excluding soldiers with an existing MDE at pre-deployment or no follow-up MDE data (resulting N=4,182). The sample was predominantly (95%) male and younger than 30 years old (mean=26.0, SD=5.9). At baseline, soldiers tended to report high trait resilience scores (mean=15.1, SD=4.0, max=20) and perceived their units as relatively cohesive (mean=19.7, SD=5.4, max=27). During deployment, soldiers reported experiencing an average of 3.8 major combat-related stressors (max=13.0, SD=2.9). Within nine months of returning from deployment, 9% (N=390) met criteria for incident MDD.

### Are polygenic risk and protective factors associated with incident MDD?

Polygenic risk showed a dose-response relationship with incident MDD. Compared to soldiers at low polygenic risk, odds for incident MDD were highest in soldiers at high polygenic risk (adjusted odds ratio [aOR]=1.52, 95% confidence interval [CI] = 1.10-2.17, p=.01) and more modestly increased in those at intermediate polygenic risk (aOR=1.33, 95% CI = 0.99-1.79, p=.06) (Figure 1a; full model results in *Supplementary Table S2B*).

**Figure 1.**
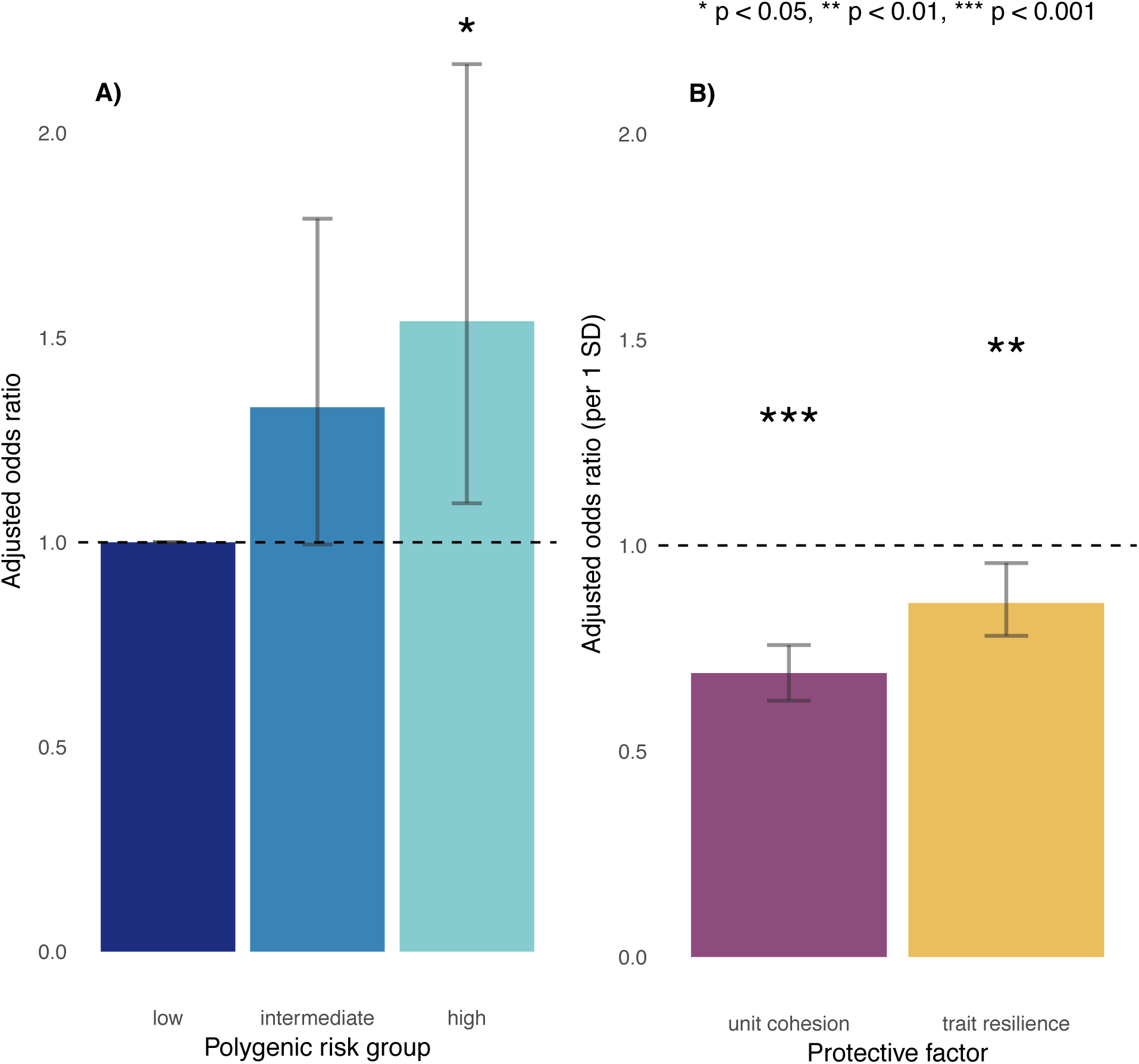
Main effects of A) polygenic risk and B) protective factors on incident MDD.

Soldiers who reported stronger unit cohesion at baseline had reduced odds for incident MDD (aOR=.69, 95% CI = 0.62-0.76, p<.0001), as did those who reported higher trait resilience (aOR=0.86, 95% CI = 0.78-0.96, p=.005 (Figure 1b). Of note, unit cohesion (r=-0.03, p=0.10) and trait resilience (r=-0.01, p=0.62) were not significantly correlated with polygenic risk, ruling out the possibility that these factors were largely determined by genetic vulnerability for MDD.

### Across the spectrum of polygenic risk, what are the effects of trait resilience and unit cohesion on incident MDD?

Across all strata of polygenic risk, soldiers who reported stronger unit cohesion had lower odds for incident MDD (*low:* aOR=0.64, 95% CI = 0.50-0.82, p=.0004; *intermediate*: aOR=0.68, 95% CI = 0.60-0.77, p<.00001; *high*: aOR=0.76, 95% CI 0.62-0.94, p=.01). Notably, even among those at highest polygenic risk, unit cohesion was associated with reduced incidence of post-deployment MDD (Figure 2a). Trait resilience showed similar but marginal effects for incident MDD across strata of polygenic risk (*low*: aOR=0.80, 95% CI = 0.62-1.04, p=.09; *intermediate*: aOR=0.90, 95% CI = 0.79-1.03, p=.13; *high*: aOR=0.83, 95% CI = 0.67-1.02, p=.08) (Figure 2b). Follow-up regressions confirmed independent and non-interactive effects of polygenic risk and unit cohesion—but not trait resilience—on incident MDD (*Supplementary Tables S2C and S2D*).

**Figure 2.**
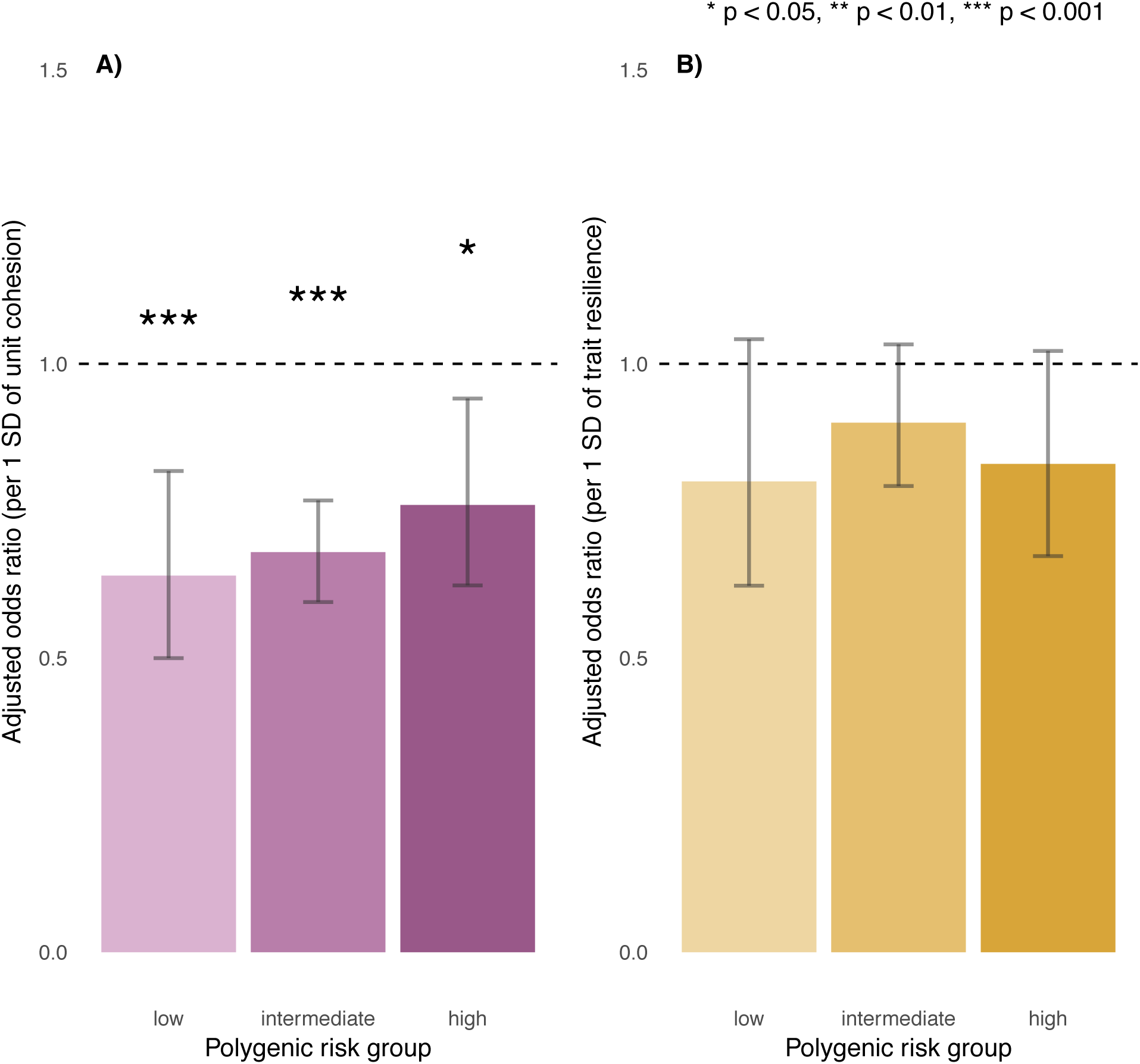
Effects of A) unit cohesion and B) trait resilience on incident MDD, stratified by polygenic risk.

### Does unit cohesion also protect against MDD after exposure to high environmental risk?

To further explore the protective effect of unit cohesion, we stratified soldiers by combat stress exposure: low (quintile 1), intermediate (quintiles 2-4), and high (quintile 5), which was itself a risk factor for incident MDD (aOR=1.46, 95% CI = 1.31-1.62, p<.00001). Unit cohesion remained protective against incident MDD across all levels of combat stress exposure (*low*: aOR=0.69, 95% CI = 0.56-0.85, p=.0005; *intermediate*: aOR=0.72, 95% CI = 0.62-0.84, p=.00002; *high*: aOR=0.63, 95% CI = 0.53-0.76, p<.00001), even for those who reported high levels of combat stress (Figure 3). Follow-up regression confirmed independent and non-interactive effects of high polygenic risk (aOR=1.48, 95% CI = 1.05-2.10, p=.03), unit cohesion (aOR=0.68, 95% CI = 0.61-0.75, p<.00001), and combat stress exposure (aOR=1.47, 95% CI = 1.32-1.63, p<.00001) on incident MDD (*Supplementary Table S2E*).

**Figure 3.**
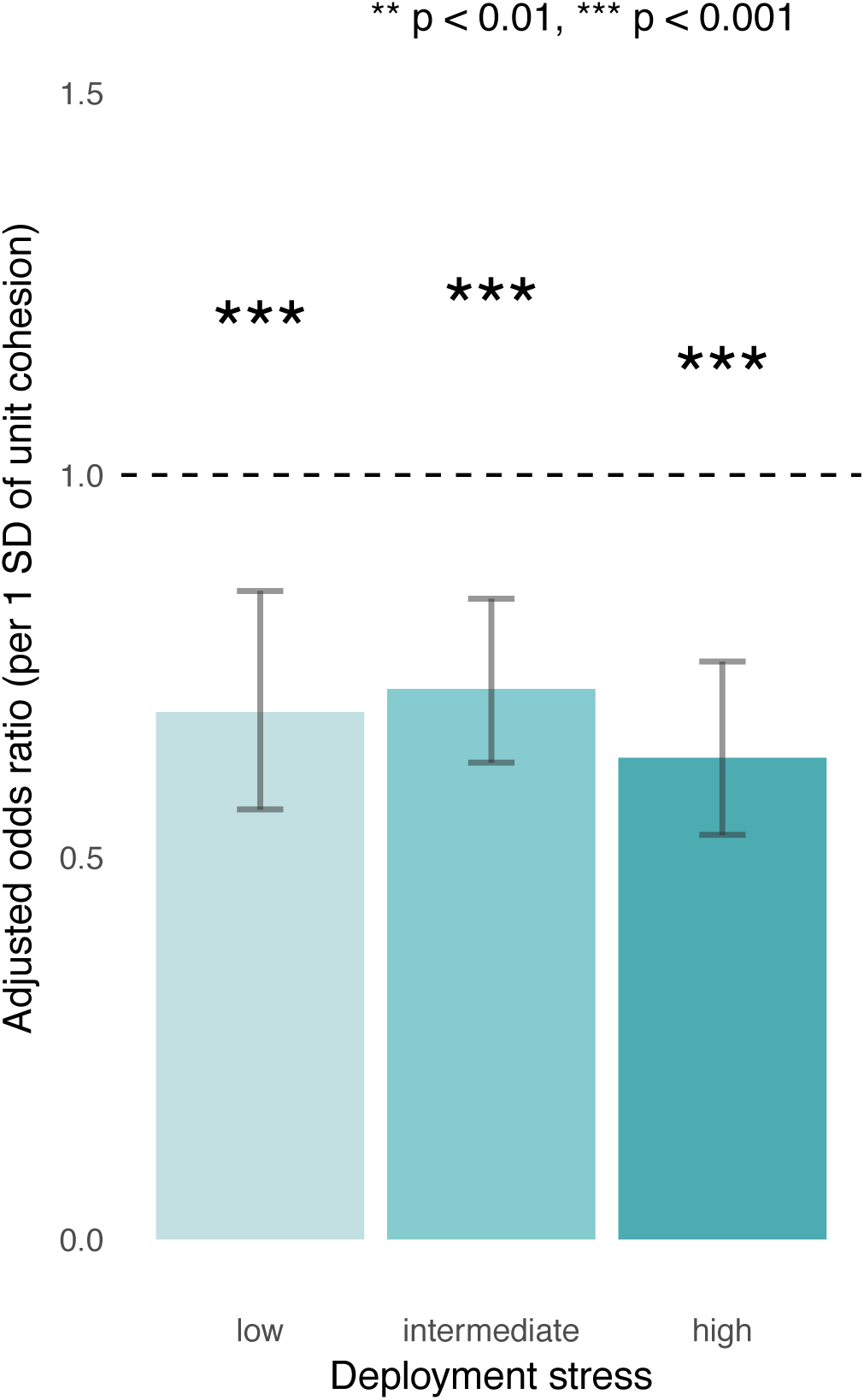
Effects of unit cohesion on incident MDD across levels of combat stress exposure.

## Discussion

In this prospective study of US Army soldiers followed across their deployment to Afghanistan, we observed that polygenic risk was associated with incident MDD following deployment; however, an extrinsic protective factor—unit cohesion—showed robust protective effects for incident MDD even among soldiers at highest polygenic or environmental risk.

Our work makes a number of contributions to the literature. First, we capitalize on a unique, large-scale, prospectively studied cohort of individuals for whom both genomic data and rich phenotyping (including exposure to a defined class of stressors) are available, to answer key questions about risk and protective factors for MDD. While genetic and environmental factors are known to influence depression, we demonstrate for the first time that polygenic risk is prospectively associated with new-onset MDD. Second, we draw on the latest and largest GWAS of MDD (10) for polygenic scoring, and validate the resulting PRS as a diathesis for MDD. Specifically, we show a dose-response relationship between polygenic risk and incident MDD following combat deployment, with a 52% increase in relative odds between soldiers in the top and bottom quintiles of polygenic risk. Thus, while PRS still account for modest variation in psychiatric traits, our main effect analyses indicate they can meaningfully explain increased risk for depression in our sample.

Third, we provide novel evidence that strong unit cohesion prior to deployment may buffer psychiatric risk regardless of underlying genetic susceptibility. Protective effects in the presence of high polygenic risk have been shown in cardiology (27) and we now apply this framework in psychiatry. While previous research has identified unit cohesion as a protective factor for mental health following deployment, most studies have been cross-sectional (16,17,28–32) and ours represents at least a four-fold increase in scale compared to existing prospective studies of unit cohesion and mental health (33,34), in addition to being the first to integrate genetic data. Fourth, we corroborate prior evidence that unit cohesion is associated with reduced risk for incident MDD despite high levels of combat stress exposure (28,31–34) and extend this to show that pre-deployment unit cohesion, combat stress exposure, and genetic susceptibility additively, and to some extent orthogonally, influence risk for incident MDD. This suggests that unit cohesion may be widely beneficial for soldiers despite genetic or environmental risk.

Unit cohesion has been conceptualized as a multi-faceted construct (35), including horizontal cohesion (e.g., perceived support from fellow soldiers, sense of bonding and camaraderie between soldiers, trust and reliance on fellow soldiers) and vertical cohesion (e.g., respect and appreciation from unit leaders; clear communication with unit leaders) (36). Our measure tapped into both aspects of unit cohesion, particularly respect and support between soldiers and with their leaders. Given inevitable stressors encountered during deployment, feeling comfortable seeking help and/or raising concerns may facilitate better coping than self-directed efforts to regulate stress (37). Moreover, strengthening such dimensions of unit cohesion is putatively actionable (38,39)—by, for example, providing leadership skills training, facilitating regular team-based interactions between soldiers during training, and keeping units operationally intact across training and deployment—though interventions remain to be rigorously tested.

Our study has several limitations. First, our analyses focused on two protective factors as exemplars of intrinsic versus extrinsic pre-deployment features, and have not tested a comprehensive set of protective factors that could reduce risk for MDD. Second, our construct of unit cohesion was measured at the individual level and may thus capture both extrinsic factors (e.g., quality of relationships, unit culture) as well as intrinsic factors that influence soldiers’ perceptions of unit cohesion (e.g., agreeableness, current distress). Future studies could utilize unit-level average cohesion scores to circumvent reporting bias and better isolate unit cohesion as an exogenous risk factor. However, individual reports may better capture the soldier’s own experience within the unit. Third, in order to establish incident MDD, we restricted our sample so that no participant met criteria for a 30-day MDE shortly before deployment. However, we did not exclude subthreshold symptoms; lifetime MDD in partial or full remission at the baseline assessment; and/or other comorbid psychopathology; and it is possible that such factors would have contributed to predicting post-deployment MDD. Fourth, our sample was primarily male and was limited to individuals of European ancestry (to maximize the power of the PRS which was based on GWAS of European ancestry subjects). Thus, our results may not generalize beyond Army populations or to female or individuals of non-European ancestry.

In conclusion, our findings support a role for both genetic and environmental factors in influencing psychiatric risk in soldiers across combat deployment. In this prospective inquiry, we showed that soldiers who experienced strong unit cohesion shortly before deployment were at reduced risk for incident MDD following deployment, regardless of their genetic susceptibility. This study illustrates the potential of protective factors to buffer psychiatric risk following exposure to stressful events. Importantly, potentially actionable factors such as group cohesion and social support may protect against depression even among those most genetically susceptible, and represent promising targets for promoting resilience in at-risk populations.

## Funding, Acknowledgements, and Conflicts of Interest

### Funding

Army STARRS was sponsored by the Department of the Army and funded under cooperative agreement number U01MH087981 (2009-2015) with the National Institutes of Health, National Institute of Mental Health (NIH/NIMH). Subsequently, STARRS-LS was sponsored and funded by the Department of Defense (USUHS grant number HU0001-15-2-0004). The contents are solely the responsibility of the authors and do not necessarily represent the views of the Department of Health and Human Services, NIMH, the Department of the Army, or the Department of Defense. Dr. Choi was supported in part by a NIMH T32 Training Fellowship (T32MH017119). Dr. Smoller is a Tepper Family MGH Research Scholar and supported in part by the Demarest Lloyd, Jr, Foundation and NIH grant K24MH094614.

### The STARRS Team

#### Co-Principal Investigators

Robert J. Ursano, MD (Uniformed Services University of the Health Sciences) and Murray B. Stein, MD, MPH (University of California San Diego and VA San Diego Healthcare System)

#### Site Principal Investigators

Steven Heeringa, PhD (University of Michigan), James Wagner, PhD (University of Michigan) and Ronald C. Kessler, PhD (Harvard Medical School) Army liaison/consultant: Kenneth Cox, MD, MPH (USAPHC (Provisional))

#### Other team members

Pablo A. Aliaga, MS (Uniformed Services University of the Health Sciences); COL David M. Benedek, MD (Uniformed Services University of the Health Sciences); Susan Borja, PhD (NIMH); Tianxi Cai, ScD (Harvard School of Public Health); Laura Campbell-Sills, PhD (University of California San Diego); Carol S. Fullerton, PhD (Uniformed Services University of the Health Sciences); Nancy Gebler, MA (University of Michigan); Robert K. Gifford, PhD (Uniformed Services University of the Health Sciences); Paul E. Hurwitz, MPH (Uniformed Services University of the Health Sciences); Kevin Jensen, PhD (Yale University); Kristen Jepsen, PhD (University of California San Diego); Tzu-Cheg Kao, PhD (Uniformed Services University of the Health Sciences); Lisa Lewandowski-Romps, PhD (University of Michigan); Holly Herberman Mash, PhD (Uniformed Services University of the Health Sciences); James E. McCarroll, PhD, MPH (Uniformed Services University of the Health Sciences); Colter Mitchell, PhD (University of Michigan); James A. Naifeh, PhD (Uniformed Services University of the Health Sciences); Tsz Hin Hinz Ng, MPH (Uniformed Services University of the Health Sciences); Caroline Nievergelt, PhD (University of California San Diego); Nancy A. Sampson, BA (Harvard Medical School); CDR Patcho Santiago, MD, MPH (Uniformed Services University of the Health Sciences); Ronen Segman, MD (Hadassah University Hospital, Israel); Alan M. Zaslavsky, PhD (Harvard Medical School); and Lei Zhang, MD (Uniformed Services University of the Health Sciences).

